# Epitranscriptomics m^6^A analyses reveal distinct m^6^A marks under tumor necrosis factor α (TNF-α)-induced apoptotic conditions in HeLa cells

**DOI:** 10.1101/2023.05.20.541583

**Authors:** Azime Akçaöz-Alasar, Özge Tüncel, Buket Sağlam, Yasemin Gazaloğlu, Melis Atbinek, Umut Cagiral, Evin Iscan, Gunes Ozhan, Bünyamin Akgül

## Abstract

TNF-α is a ligand that induces both intrinsic and extrinsic apoptotic pathways in HeLa cells by modulating complex gene regulatory mechanisms. However, the full spectrum of TNF-α-modulated epitranscriptomic m^6^A marks is unknown. We employed a genomewide approach to examine the extent of m^6^A RNA modifications under TNF-α-modulated apoptotic conditions in HeLa cells. miCLIP-seq analyses revealed a plethora of m^6^A marks on 632 target mRNAs with an enrichment on 99 mRNAs associated with apoptosis. Interestingly, the m^6^A RNA modification patterns were quite different under cisplatin- and TNF-α-mediated apoptotic conditions. We then examined the abundance and translational efficiencies of several mRNAs under METTL3 knockdown and/or TNF-α treatment conditions. Our analyses showed changes in the translational efficiency of *TP53INP1* mRNA based on the polysome profile analyses. Additionally, TP53INP1 protein amount was modulated by METTL3 knockdown upon TNF-α treatment but not CP treatment, suggesting the existence of a pathway-specific METTL3-TP53INP1 axis. Congruently, METLL3 knockdown sensitized HeLa cells to TNF-α-mediated apoptosis, which was also validated in a zebrafish larval xenograft model. These results suggest that apoptotic pathway-specific m^6^A methylation marks exist in cells and TNF-α-METTL3-TP53INP1 axis modulates TNF-α-mediated apoptosis in HeLa cells.

## Introduction

Extracellular and intracellular signals play a fundamental role in the critical balance between cell death and survival that is required to maintain homeostasis throughout development and to prevent diseases such as cancer or neurodegenerative diseases (Flusberg and Sorger 2015; Gudipaty et al. 2018). Interestingly, some signals may lead to opposite cellular effects under different cellular conditions in a cell specific manner. For example, while cisplatin (CP) typically triggers the intrinsic apoptotic pathway, the binding of tumour necrosis factor-α (TNF-α) ligand to its receptor may lead to cell death or survival in a cell-specific manner (Wajant, Pfizenmaier, and Scheurich 2003). TNF-α is a transmembrane protein released as a soluble ligand upon proteolytic cleavage (Black et al. 1997). Its interaction with TNF receptors (TNFRs) may lead to apoptosis by activating the extrinsic pathway (L. Wang, Du, and Wang 2008). Most molecular mechanisms underlying transcriptional and posttranscriptional regulation of survival and cell death have been well-documented and targeted for various therapeutic purposes (Sedger and McDermott 2014; Tüncel et al. 2022). However, recent studies point to the importance of epitranscriptomics as a new layer of gene regulation in modulation of survival and cell death that might serve as novel therapeutic targets (Akçaöz and Akgül 2022).

The *N*^6^-methyladenosine (m^6^A) modification constitutes the most abundant epitranscriptomics mark in cellular RNAs being present in 0.1-0.4% of all adenosines (Wei, Gershowitz, and Moss 1975). A specific set of proteins called writers, readers and erasers perform deposition, recognition and removal of m^6^A RNA marks, respectively (Zaccara, Ries, and Jaffrey 2019). The reversible nature of cell- and tissue-specific m^6^A RNA marks play an instrumental role in modulating the transcriptional or posttranscriptional fate of mRNAs (Yue, Liu, and He 2015). Recent studies clearly exhibit that perturbations in the expression of writers, readers or erasers are associated with survival and cell death (Akçaöz and Akgül 2022). We have recently reported that the cancer chemotherapeutic drug CP modulates the differential m^6^A RNA methylation of numerous mRNAs (Alasar et al. 2022). Interestingly, CP-mediated apoptosis appears to be influenced by METTL3-dependent translational regulation of PMAIP1, which is a downstream effector of p53 (Oda et al. 2000).

In contrast to CP, TNF-α is a ligand that primarily induces the extrinsic apoptotic pathway (L. Wang, Du, and Wang 2008). Recent reports have shown that the cellular effects of TNF-α may be modulated by m^6^A writers, readers or erasers. For example, directional migration of mesenchymal stem cells in ankylosing spondylitis is mediated by m^6^A modification of engulfment and cell motility 1 (*ELMO1*) in a TNF-α-dependent manner as revealed by methyltransferase like 14 (METTL14) knockdown (Xie et al. 2021). Similarly, sweet-gland differentiation of mesenchymal stromal cells by TNF-α involves fat mass and obesity-associated protein (FTO)-mediated m^6^A-demethylation of *NANOG* mRNA (Y. Wang et al. 2020). Demethylase α-ketoglutarate-dependent dioxygenase ALKB homolog 5 (ALKBH5) has been shown to promote proliferation and to inhibit apoptosis in TNF-α-treated human umbilical vein endothelial cells (Xiaoshan Zhang et al. 2022). Although these studies clearly suggest the significance of m^6^A epitranscriptional modulation of TNF-α signalling, the full spectrum of m^6^A RNA marks modulated by TNF-α is unknown.

In this study, we exploited the miCLIP-seq approach to uncover the extent of m^6^A RNA methylation in TNF-α-treated HeLa cells. Our results have shown that CP- and TNF-α-induced apoptotic cells display quite a distinct m^6^A RNA profile. Of 632 number of mRNAs differentially m^6^A-methylated, 99 mRNAs were associated with apoptotic signalling. qRT-PCR and polysome analyses identified tumor protein 53 inducible nuclear protein 1 (TP53INP1) as a target of METTL3. Interestingly METTL3 knockdown further sensitized HeLa cells to TNF-α-mediated apoptosis by enhancing its translation efficiency whereas CP did not have any detectable effect on the translational efficiency of the *TP53INP1* transcript in METTL3-knockdown HeLa cells.

## Materials and Methods

### Cell Culture, Drug Treatments and Analysis of Apoptosis

HeLa and ME-180 cervical cancer cell lines were purchased from DSMZ GmbH (Germany) and ATCC (United States), respectively. HeLa and ME-180 cells were cultured with RPMI-1640 (Gibco) and McCoy’s 5A (Lonza, Switzerland), respectively, supplemented with 10% FBS (Gibco) in a humidified atmosphere of 37°C and 5% CO_2_. HeLa cells were treated with TNF-α recombinant human protein (Biolegend) at a concentration of 75 ng/ml for 24 h in combination with cycloheximide (CHX) (10 µg/ml) (Applichem) to trigger the extrinsic apoptotic pathway. CHX treatment without TNF-α protein was used as negative control. To induce the intrinsic apoptotic pathway, cells were treated with 80 µM CP (Santa Cruz Biotechnology) and 0.1% (*v/v*) DMSO for 16 h (Yaylak, Erdogan, and Akgul 2019).

In transfected cells, the reduced dose of TNF-α and CP were used to compensate for the apoptotic effect of the transfection reagent. Accordingly, transfected cells were treated with 2.5 µg/ml CHX and 37.5 ng/ml TNF-α recombinant human protein or 40 µM CP. Subsequently, cells were trypsinized by trypsin-EDTA (0.25%) (Gibco), centrifuged at 1000 rpm for 5 min and washed with 1X cold PBS (Gibco). The resulting pellet was suspended with 1X Annexin binding buffer (Becton Dickinson), followed by staining with AnnexinV-FITC (Biolegend, USA) and 7AAD (Biolegend). After 15 min of incubation at room temperature (RT) in the dark, apoptosis rate was measured by flow cytometry (BD FACSCanto) as previously published (Yaylak, Erdogan, and Akgul 2019).

### Cell Transfection

The METTL3 overexpression plasmid pcDNA3.1(+)-METTL3 was described previously (Alasar et al. 2022). HeLa cells were transfected with pcDNA3.1(+)-METTL3 or pcDNA3.1(+) using Fugene HD transfection reagent (Promega) according to the manufacturer’s protocol. Briefly, 75,000 cells/well were seeded into 6-well plates (Sarstedt), and transfection was performed at 80% confluence. Cells were transfected with 1.5 µg plasmid DNA/well using the transfection reagent at a ratio of 1:3. The culture medium was changed 1 h post-transfection, and cells were harvested 48 h post-transfection.

For knockdown experiments, a total of 75,000 cells/well HeLa cells were seeded on 10 cm dishes (Sarstedt) overnight, and were transfected with 25 nM non-target pool siRNA (si-NC) or METTL3 siRNA (si-METTL3) (Dharmacon) using DharmaFECT transfection reagent at a ratio of 2:1 (*v/v*) in 800 μL serum-free medium. Transfected cells were incubated for 72 h prior to harvesting for subsequent experiments.

### Total RNA Isolation and qPCR

TRIzol^TM^ reagent (Invitrogen) was used for total RNA isolation according to the manufacturer’s instructions. cDNA was prepared with 2 µg of total RNA using RevertAid first strand cDNA synthesis kit (Thermo Fisher Scientific) according to the manufacturer’s protocol. qPCR reactions were set up with GoTaq® qPCR Master Mix (Promega) using diluted cDNA (5 ng/µl). qPCR primers were listed in Table 1. All reactions were run in a Rotor-Gene Q machine (Qiagen) as follows: initial denaturation at 95°C for 2 min followed by 45 cycles of denaturation at 95°C for 15 s and annealing at 60°C for 1 min. GAPDH was used for normalization.

**Table 1.**
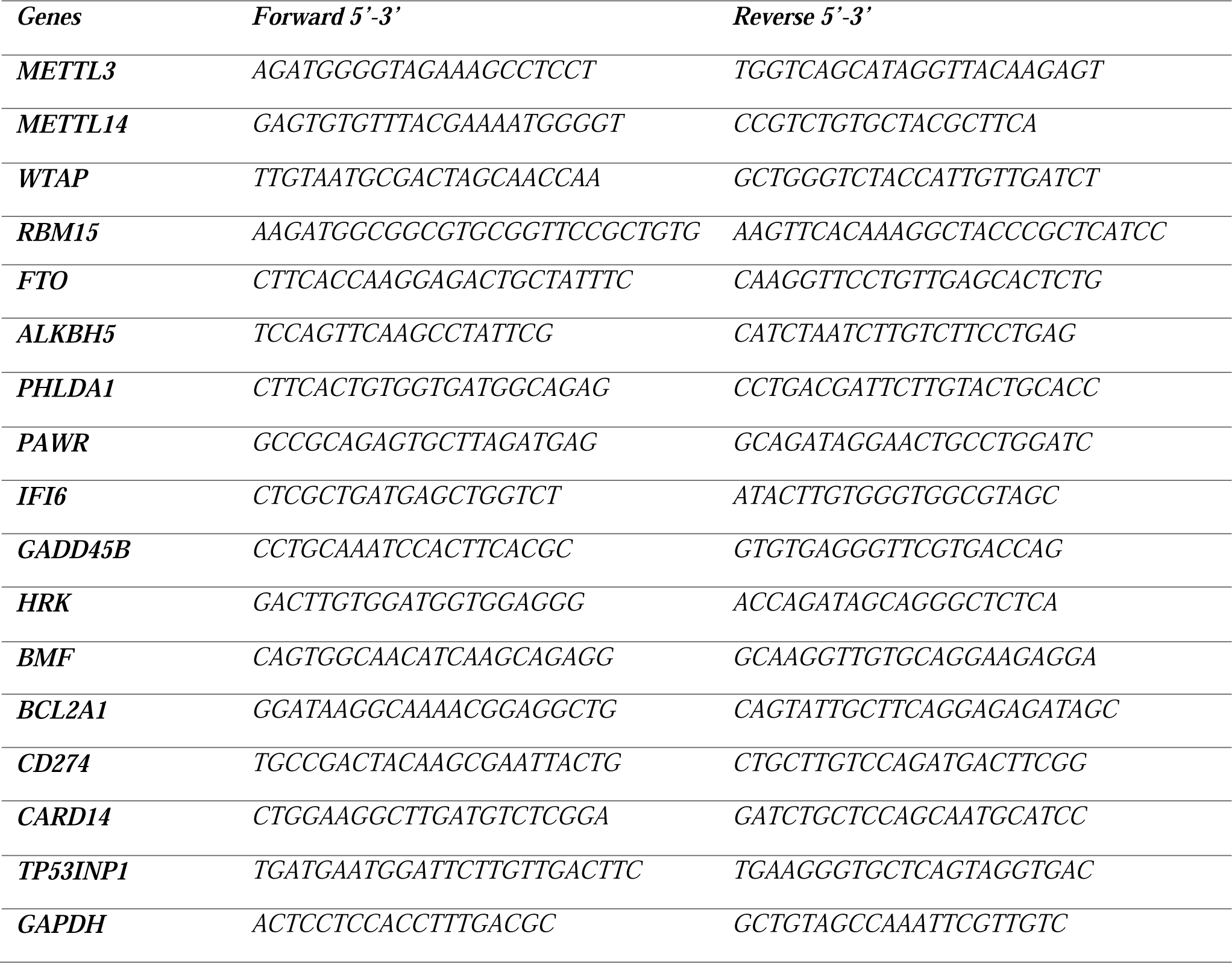
The list of primers used in qPCR analyses.

### M^6^ A-eCLIP-seq and Bioinformatic Analyses

Total RNAs were extracted from TNF-α-treated HeLa cells by TRIzol. The samples were subjected to m^6^A-eCLIP by Eclipse BioInnovations Inc, San Diego CA as described in their user guide. The data were deposited to Genome Expression Omnibus with the accession number GSE223209. Data analysis was performed by Eclipse BioInnovations using their standard m^6^A-eCLIP analysis pipeline as described in Alasar et al. 2022. Differentially methylated transcripts were analyzed by Gene Ontology enrichment tool. Heatmaps were generated using m^6^A-eCLIP-seq fold change data. All analyses were performed using R (R Core Team, 2020), RStudio (Rstudio Team, 2020), and the “pheatmap” package 1.0.12. (Kolde, 2019). Treatments and genes were clustered, and the maps were scaled by rows.

### Western Blotting

Total proteins were extracted by using protease inhibitor and RIPA buffer (CST). 25 µg of total protein extract was fractionated on a 5–10% sodium dodecyl sulphate (SDS) polyacrylamide gel. The PVDF membranes with transferred proteins were blocked for 1 h at RT in Tris-buffered saline (TBS) containing 0.2% (v/v) Tween-20 and 5% nonfat dry milk. The membranes were incubated overnight with primary monoclonal antibodies against METTL3 (Rabbit mAb #96391, CST), METTL14 (Rabbit mAb #51104S, CST), FTO (Rabbit mAb #31687S, CST), RBM15 (#25261S, CST), TP53INP1 (ab202026, Abcam), Caspase 8 (Mouse mAb #9746, CST), Caspase 9 (Mouse mAb #9508, CST), Caspase 3 (Rabbit mAb #14220, CST) and ß-actin (Rabbit mAb #4970, CST) at 4°C. Then, membranes were incubated with secondary antibodies (1:2000) in TBS-T with 5% nonfat dry milk for 1Uh at RT and washed 3 times for 5 min with TBS-T. Membranes were incubated in HRP, and chemiluminescent signals were quantified by the ImageJ Software. β–actin was used as the loading control.

### Polysome Analyses

Polysome profiling was carried out as described previously (Göktaş et al. 2017). Cell pellets were resuspended in the homogenization buffer [100 mM NaCl, 10 mM MgCl_2_, 30 mM Tris-HCl (pH 7), 1% Triton X-100, 1% NaDOC, 100 µg/mL CHX (Applichem) and 30 U/mL RNase Inhibitor (Promega)]. Cells were disrupted with a 26 G syringe and homogenized by passing the lysate through the needle at least 15 times to obtain a homogeneous lysate. The cell lysates were first incubated on ice for 8 min followed by centrifugation at 12,000 *g* at 4°C for 8 min. The supernatants were then layered over 5-70% (w/v) sucrose gradients prepared with the homogenization buffer except for CHX, NaDOC and Triton X-100 and centrifuged at 27,000 rpm for 2 h 55 min at 4°C in a Beckman SW28 rotor. Fractions were pooled into messenger ribonucleoprotein (mRNP), monosomal, light and heavy polysomal sub-groups based on absorbance values at 254 nm. Total RNAs were phenol-extracted from the fractions by using phenol-chloroform-isoamyl alcohol solution (25:24:1) as previously described (Alasar et al. 2022).

### Zebrafish Larval Xenografts

2 days post-fertilization (dpf) *casper* (*roy -/-; nacre-/-)* zebrafish larvae were dechorionated with 0.1 mg/mL pronase (Sigma-Aldrich, MO, USA) solution for 5-7 min at 28°C, anesthetized with 1 mg/mL Tricaine in E3 embryo medium and transferred to a microinjection plate prepared with 3% agarose in E3 medium. Microinjection was performed using borosilicate glass capillaries (4 inches, OD 1.0 mm, World Precision Instruments, FL, USA). 200-250 HeLa cells transfected with si-NC or si-METTL3 at a concentration of 25 nM were injected directly into the middle of the yolk sac of the larva, preventing damage to the duct of Cuvier. Larvae were treated with 10 ng/ml TNF-α and 2.5 µg/ml CHX and incubated at 34°C in fresh E3 medium for 24 h. The next day, 1-day post-injection (dpi), larval xenografts were processed for immunofluorescence staining.

### Whole Mount Immunofluorescence Staining, Confocal Imaging and Quantification of Zebrafish Larvae

The immunostaining procedure previously described by Martinez-Lopez et al. was followed with several modifications (Martinez-Lopez et al. 2021). Larvae were fixed in 4% paraformaldehyde (PFA) in 1X PBS overnight at 4 °C, washed with 1x PDT (1x PBST, 0.3% Triton-X, 1% DMSO) and permeabilized with ice-cold acetone. The larvae were blocked for 2 h in PBDX GS blocking buffer (%10 bovine serum albumin, %1 DMSO, 0.3% Triton-X, 15 µL/1 mL goat serum) and incubated with the primary antibody, rabbit anti-cleaved-caspase-3 (1:200, 5A1E, Cell Signaling Technology, MA, USA), in PBS/0.1% Triton at 4 °C overnight. The next day, larvae were washed with PBS/0.1% Triton, incubated with the secondary antibody, Fluorescein (FITC) AffiniPure donkey anti-rabbit IgG (1:200, 711-096-152, Jackson Immunoresearch Laboratories, PA, USA), at RT for 2 h, refixed in 4% PFA at RT for 20 min, and washed with PBS/0.1% Triton. Following nuclear staining with 4′,6-diamidino-2-phenylindole (DAPI; 4083S, Cell Signaling Technology, MA, USA), larvae were mounted in 80% glycerol between two coverslips and imaged using fluorescence confocal microscopy. Confocal images were recorded with a 25x objective lens. Image processing and quantifications were performed using FIJI/ImageJ software by following the previously published protocol (Martinez-Lopez et al. 2021). To quantify apoptosis, cleaved-caspase-3 positive cells were counted using the counter plugin and divided by the number of DiL+, DAPI+ nuclei in each corresponding slice.

## Results

### TNF-α Perturbs the Expression of the m^6^A Methylation Apparatus

We previously reported that CP, a well-known inducer of intrinsic apoptotic pathways, modulates apoptosis by differential m^6^A RNA methylation of a select set of transcripts, *PMAIP1* being one of those targets (Alasar et al. 2022). Since intrinsic and extrinsic apoptotic pathways are regulated through different transcriptional and post-transcriptional mechanisms (L. Wang, Du, and Wang 2008), we hypothesized that there should be m^6^A RNA methylation marks specific to extrinsic apoptotic pathways. To uncover the extrinsic pathway-specific m^6^A RNA methylome, we used TNF-α to induce the extrinsic apoptotic pathways in HeLa cells. CHX was coupled to TNF-α to switch the inflammatory response of TNF-α to apoptosis (L. Wang, Du, and Wang 2008). TNF-α at a concentration of 75 ng/mL triggered an early apoptotic rate of 35.1% compared to 10.2% in the control CHX-treated HeLa cells (Figure 1A-B). We compared the extent of activation of intrinsic and extrinsic pathways by examining the cleavage patterns of caspases in TNF-α- and CP-treated cells. As expectedly, CP caused primarily activation of caspase-9 with a nearly undetectable amount of cleaved caspase-8 (Figure 1C). On the other hand, we detected a prominent cleavage of caspase-8 in TNF-α-treated cells, which also activated the intrinsic pathway as revealed by the cleavage of caspase-9 (Figure 1C). Treatment with both inducers triggered caspase-3 cleavage, a biochemical indicator of activation of apoptosis.

**Figure 1.**
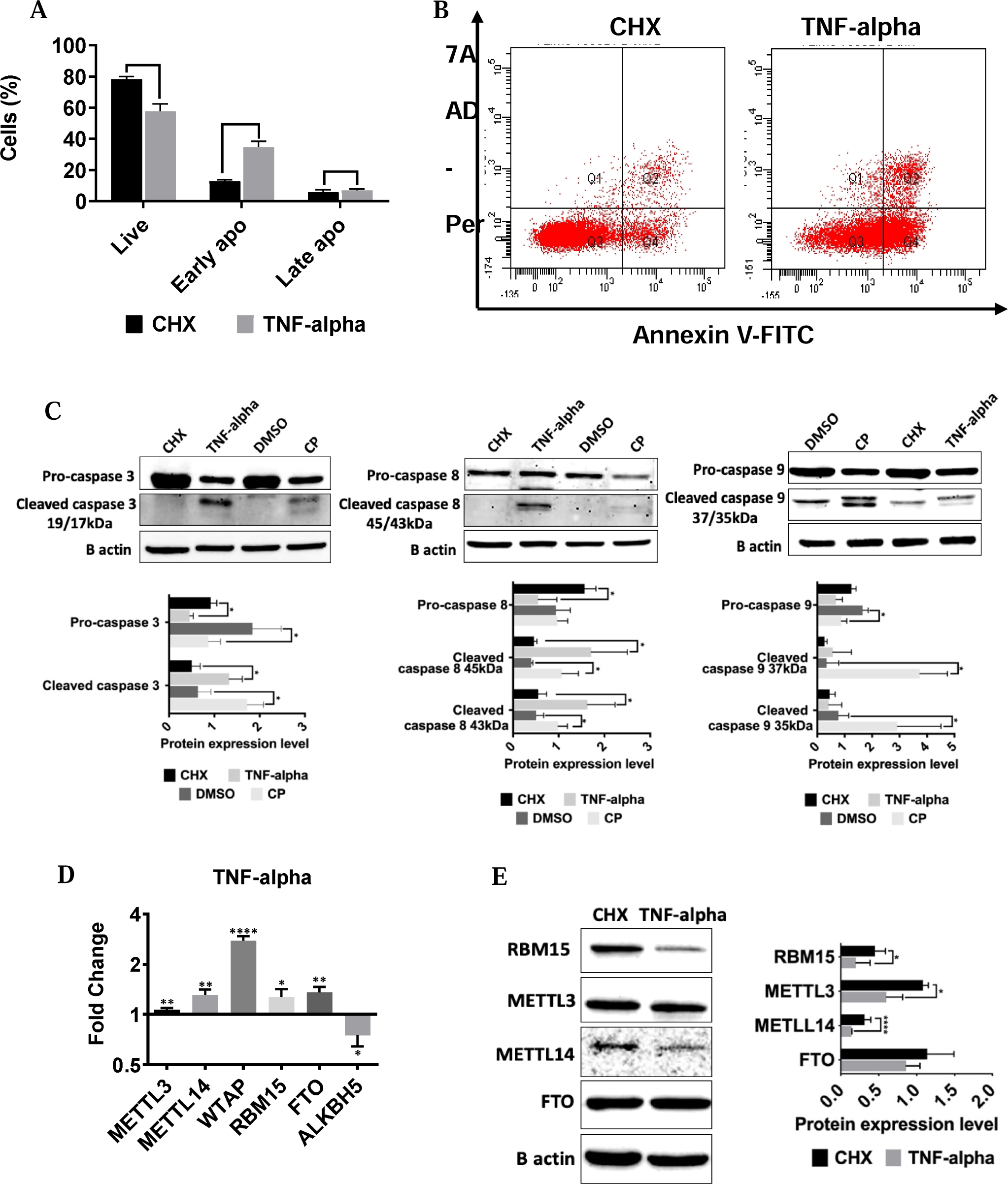
TNF-alpha treatment affects the expression of m^6^A enzymes in HeLa cells. HeLa cells were treated with 75 ng/ml TNF-α and 10 µg/ml CHX for 24h. The cell populations were quantified by Flow cytometry using Annexin V and 7AAD. Error bars represent meanU±USD of three independent experiments with at least 10,000 cells counted per treatment (unpaired, two-tailed t-test). A. The percentage of live, early and late apoptotic cells. B. Dot-blot analysis by flow cytometry after staining with Annexin V-PE and 7AAD. C. Immunoblot analyses of caspase -3, -8, and -9 in TNF-α-(75 ng/ml, 24h) and CP-treated (80 µM, 16h) HeLa cells. β-actin was used as a loading control. D. qPCR analyses of the expression levels of m^6^A writers and erasers in TNF-α -treated HeLa cells. Data were normalized to GAPDH. E. Western blot analyses of m^6^A writers and erasers in TNF-α-treated HeLa cells. Data were normalized to ß-actin. CHX, cycloheximide. Error bars represent meanU±USD of three independent experiments. Two-tailed Student’s t test was performed to determine the statistical significance among groups. *: p≤0.05, **: p≤0.01, ****: p≤0.0001

To examine whether the m^6^A RNA methylation machinery is modulated by TNF-α-mediated apoptotic conditions, we measured the mRNA and protein levels of m^6^A writers and erasers. TNF-α treatment of HeLa cells led to a 2.8-fold increase in the *WTAP* transcript abundance with minor changes in the abundance of *METTL3, METTL14, RBM15, FTO* and *ALKBH5* transcripts (Figure 1D). On the other hand, we detected 2.3-, 2.2-, 1.8- and 1.3-fold decreases on the protein levels of RBM15, METTL14, METTL3 and FTO, respectively (Figure 1E).

### A Distinct Set of m^6^A RNA Marks Are Deposited Upon TNF-α and Cisplatin Treatments

It is very interesting that METTL3 and METTL14 protein amounts are downregulated upon both CP and TNF-α treatments (Alasar et al. 2022; Figure 1C). There appears to be a consistent reduction in the writer protein amounts irrespective of the apoptotic inducer. However, the differential proteolytic cleavage of caspase-8 and -9 by CP and TNF-α suggests activation of different signal transduction pathways, and different m^6^A RNA marks as a result. We hypothesized that a different set of transcripts should be m^6^A RNA methylated under TNF-α treatment conditions to exert a pathway specific apoptosis probably through the modulatory effect of unknown accessory protein(s). Thus, we performed m^6^A miCLIP-seq analyses with total RNAs isolated from TNF-α-treated HeLa cells and compared the resulting m^6^A methylome to that obtained under CP treatment conditions (Figure 2A-B; Alasar et al. 2022). TNF-α treatment in HeLa cells resulted in differential m^6^A RNA methylation of 632 transcripts with elevated methylation at 2941 positions and reduced methylation in 987 positions (Supplementary Table 1). Strikingly, the heat map constructed from differentially m^6^A methylated transcripts displayed a stimulus-specific m^6^A RNA marks (Figure 2C). When we examined the metaintron plots of up- and downregulated m^6^A RNA marks in TNF-α treated cells (Figure 2D and E, respectively), we noticed a similar enrichment based on the relative position on transcripts.

**Figure 2.**
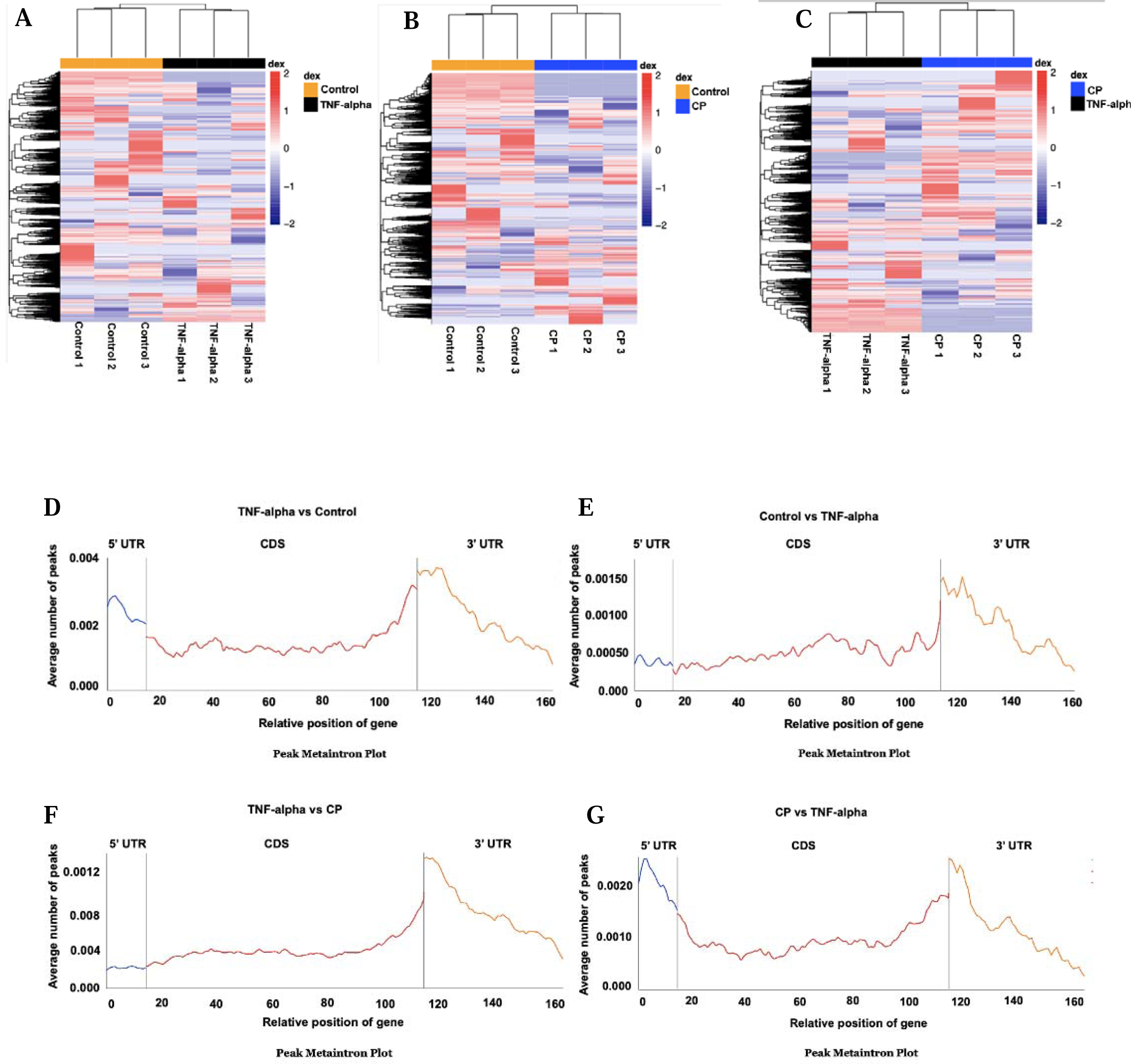
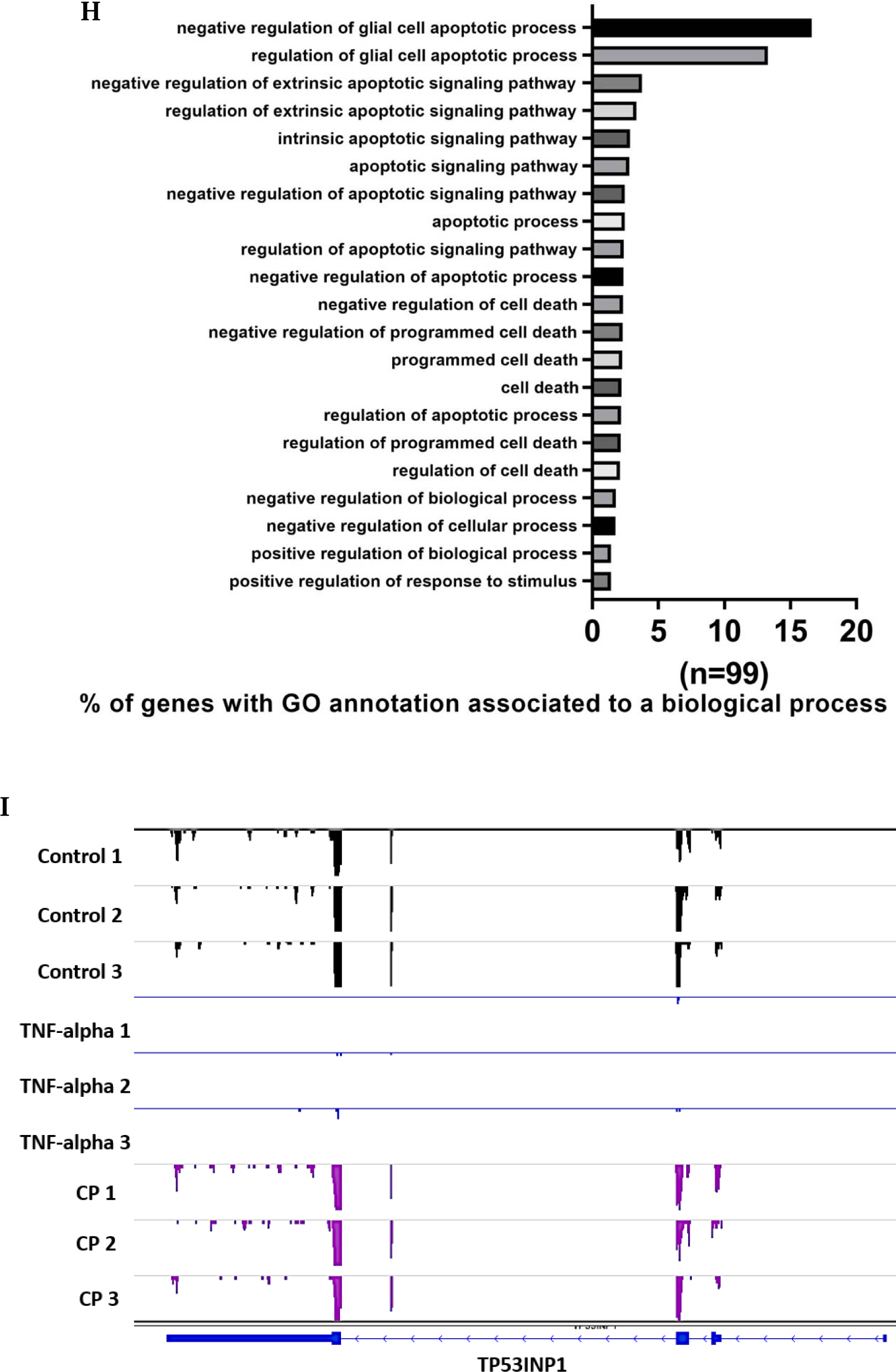
m^6^A methylome analysis of TNF-α-treated HeLa cells. miCLIP-seq was used to obtain the m^6^A methylome of TNF-alpha treated HeLa cells as outlined in Materials and Methods. A-C. Heatmap of differentially m^6^A-methylated transcripts in CP- and TNF-α-treated HeLa cells. The list of differentially m^6^A-methylated transcripts in CP-treated cells was published previously (Alasar et al. 2022). Metaintron profiles of transcripts upregulated (D) or downregulated (E) in m^6^A methylation in TNF-α-treated HeLa cells and upregulated (F) or downregulated (G) in TNF-α-induced apoptosis compared to CP-treated HeLa cells along a normalized transcript, consisting of three rescaled non-overlapping segments: 5’UTR, CDS, and 3’UTR. H. Gene Ontology analyses of differentially m^6^A-methylated genes associated with apoptosis. I. Distribution of m^6^A modification along the CDS, 3’UTR and 5’UTR of *TP53INP1* mRNA was displayed by Integrative Genomics Viewer (IGV): Control (upper panels), as a control, TNF-α (middle panels) and CP treatment (lower panels).

However, when the enrichment patterns of up- and down-regulated m^6^A RNA marks were compared between TNF-α- and CP-treated cells, we observed a different pattern (Figure 2F-G, respectively). This observation was affirmed by a similar distribution pattern on different regions of transcripts (Supplementary Figure 1). When we performed Gene Ontology (GO) analyses with the differentially methylated transcripts, we noticed enrichment in numerous biological processes, apoptotic processes being among them (Supplementary Table 2). Of 632 differentially methylated transcripts, 99 of them was associated with cell death/apoptosis (Figure 2H, Supplementary Table 3), suggesting that TNF-α-mediated apoptosis involves modulation of m^6^A RNA methylation of apoptosis-related transcripts. TNF-α-modulated increase in the m^6^A marks of one of those transcripts, *TP53INP1*, is illustrated in Figure 2I.

### METTL3 Knockdown Sensitizes HeLa Cells to TNF-α-mediated Apoptosis

We previously reported that METTL3 knockdown leads to a 5.8% elevation in the rate of early apoptosis in CP-treated Hela cells (Alasar et al. 2022), suggesting that METTL3 knockdown may sensitize HeLa cells to the activation of CP-induced intrinsic apoptotic pathways. To investigate whether this observation is specific to CP-mediated apoptosis or a more general cellular response, we examined the sensitization of HeLa cells to TNF-α-mediated apoptosis upon METTL3 knockdown. To this extent, we carried out TNF-α treatment following METTL3 knockdown (Figure 3A-B). Interestingly, the extent of sensitization was much greater under TNF-α-mediated apoptotic conditions, compared to CP treatment (Alasar et al. 2022), with the rate of early apoptotic HeLa cells increasing from 18.2% to 42% (Figure 3B). We hypothesized that TNF-α-mediated apoptosis and further sensitization of HeLa cells to apoptosis by METTL3 knockdown could be mediated by some of differentially m^6^A methylated transcripts (Figure 2). Since m^6^A methylation has been reported to modulate RNA abundance or translatability (Zaccara, Ries, and Jaffrey 2019), we selected a set of 10 apoptotic transcripts with highest differential m^6^A RNA methylation under TNF-α-mediated apoptotic conditions, to examine whether their abundance is correlated with METTL3 knockdown. To this extent, we first interrogated the transcript abundance under TNF-α treatment and METTL3 knockdown separately (Figure 3C-D). Our results showed that TNF-α treatment led to a 1.4-, 2.1-, 2.7- and 4.6-fold increase in the expression of *HRK*, *CD274*, *PHLDA1*, and *BCL2A1*, and 1.3- and 4.8-fold decrease in the expression of *PAWR* and *BMF*, respectively (Figure 3C). METTL3 knockdown resulted in a 1.4-, 1.8-, and 2.9-fold decrease in the expression of *GADD45B*, *PAWR*, and *HRK* and a 1.3- and 1.5-fold increase in the expression of *IFI6* and *PHLDA1*, respectively (Figure 3D). This observation suggests that although METTL3 knockdown does not have much of an effect on the abundance of the candidate apoptotic transcripts tested (Figure 3D), TNF-α modulates the abundance of at least *PHLDA1*, *BMF*, *BCL2A1* and *CD274* (Figure 3C). We then examined how TNF-α treatment affects the abundance of these transcripts upon METTL3 knockdown. Although we detected minor differences in the extent of differential expression of some transcripts between control (Figure 3E, CHX only) and TNF-α-treated (Figure 3F, CHX and TNF-α) cells, typically the expression patterns were quite similar.

**Figure 3.**
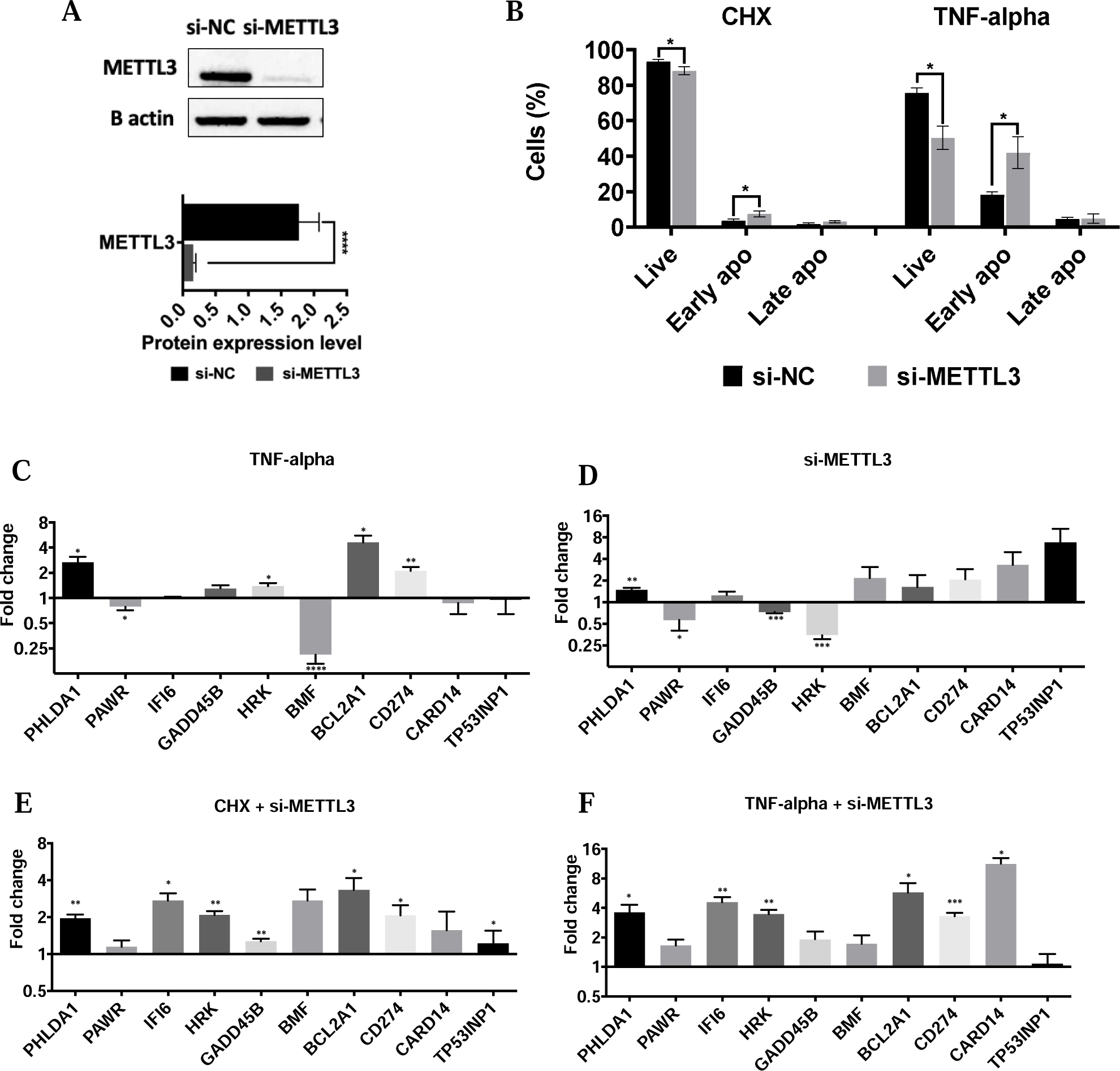
Effects of METTL3 depletion on RNA abundance. A. Western blot analysis of HeLa cells transfected with METTL3 siRNA (si-METTL3). Negative control was non-targeting pool siRNA (si-NC) and loading control was ß-actin. B. The rate of apoptosis in HeLa cells transfected with 25 nM si-METTL3 for 72h and/or incubated with 37.5 ng/ml TNF-α for 24h. Cells were stained with Annexin V and 7-AAD, and analyzed by flow cytometry. qPCR analyses of *PHLDA1*, *PAWR, IFI6, GADD45B, HRK, BMF, BCL2A1, CD274, CARD14* and *TP53INP1* 24h after TNF-alpha treatment (C), 72h after METTL3 depletion (D), CHX treatment with METTL3 knockdown (E) and TNF-alpha treatment with METTL3 knockdown. All data are representative of three independent experiments. CHX, cycloheximide. Two-tailed Student’s t test was performed to determine the statistical significance among groups. All data are presented as meanU±USD. *: p≤0.05, **: p≤0.01, ***: p≤0.001****: p≤0.0001

### TNF-α-mediated Translational Efficiency of mRNAs is Modulated by METTL3

Current studies suggest that m^6^A RNA modifications could potentially change the secondary structure of mRNAs and modulate their translational efficiencies by interfering with RNA:RNA or RNA:protein interactions (Zaccara, Ries, and Jaffrey 2019). To examine the impact of METTL3 in the translational efficiencies of candidate apoptotic transcripts under TNF-α-mediated apoptotic conditions, we aimed to quantitatively measure the association of candidate transcripts with polysomes as an indication of efficient translation (Chasse et al., 2017). To this extent, we first examined whether METTL3 knockdown, without induction of apoptosis, affects the translatability of selected transcripts. We previously reported the fractionation of HeLa cells transfected with si-NC (control) and si-METTL3 on sucrose density gradients (Alasar et al. 2002). The fractions were pooled into four major fractions based on the A_254_ readings, namely (1) mRNP fraction containing mRNAs unassociated with any ribosomes or ribosomal subunits, (2) monosome fraction harbouring mRNAs associated with a single ribosome, (3) light polysome fraction containing mRNAs loaded with a few ribosomes and (4) heavy polysome fraction that contains mRNAs associated more ribosomes. To examine the effect of METTL3 knockdown on individual candidate transcripts, we quantitatively measured the abundance of selected transcripts by qPCR analyses of total RNAs extracted from these four major fractions. Our analyses revealed that METTL3 knockdown perturbs the polysomal association of *PAWR*, *IFI6* and *TP53INP1*, the nonpolysomal (mRNP or monosome) association of *IFI6*, *GADD45B*, *HRK*, *BMF* and *BCL2A1*, without any major effects on the translation efficiency of *PHLDA1* and *CARD14* (Supplementary Figure 2).

To interrogate the polysomal association of the apoptotic candidate transcripts in the presence or absence of TNF-α in METTL3 knockdown cells, we performed polysome profiling analyses with HeLa cells as previously reported (Alasar et al. 2022). TNF-α treatment did not change the polysome profile of METTL3 knockdown cells compared to the control CHX-treated cells (Figure 4A). Although METTL3 knockdown perturbed the polysomal association of some candidates under TNF-α treatment conditions (Figure 4B-K), some others were not affected by TNF-α treatment (Supplementary Figure 3). Of all apoptotic candidates tested, the *TP53INP1* transcript stroke our attention for two reasons: (1) it is a target of p53 (Saadi, Seillier, and Carrier 2015), and (2) it is highly associated with polysomes under TNF-α treatments while it appears to be associated with mRNP and monosomes under the control CHX treatment conditions (Figure 4J-K).

**Figure 4.**
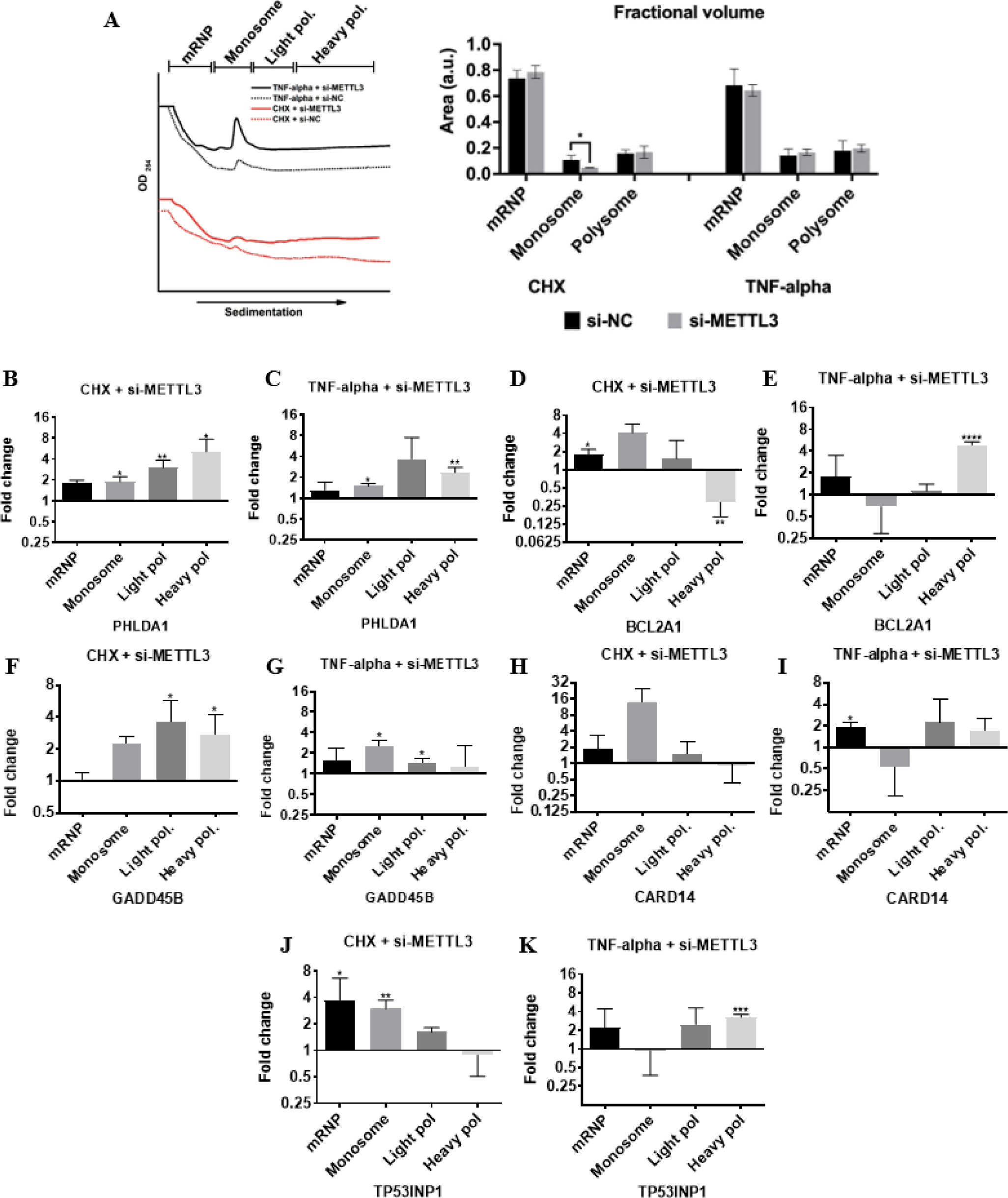
Effects of METTL3 knockdown and TNF-α on translational efficiencies of candidate mRNAs. A. Polysome profile of HeLa cells transfected with 25 nM control si-NC (dotted lines) or si-METTL3 (continuous lines) and treated with 2.5 µg/ml CHX (red lines) and 37.5 ng/ml TNF-α for 24 h (black lines). Cleared cytoplasmic cell lysates were fractionated into mRNP, monosome, light and heavy polysomes in 5−70% sucrose gradients by simultaneous detection of absorbance at 254 nm. Areas under the peak of each fraction was quantified and plotted as percentages of the total area under the full profile. mRNA abundance of *PHLDA1* (B, C), *BCL2A1* (D, E), *GADD45B* (F, G), *CARD14* (H, I) and *TP53INP1* (J, K) mRNAs in each fraction was quantified by qPCR. n = 3 biological replicates. CHX, cycloheximide. Two-tailed Student’s t test was performed to determine the statistical significance among groups. Data presented as mean ± SD, *: p≤0.05, **: p≤0.01, ***: p≤0.001****: p≤0.0001.

Although the TNF-α-mediated switch of *TP53INP1* transcripts from nonpolysomal fractions to polysomal fractions suggests that *TP53INP1* is likely to be translated more efficiently under TNF-α treatment and METTL3 knockdown conditions, it may exist in a high-molecular-weight pseudo-polysome complex (Thermann and Hentze 2007), other than polysomes, that could cause it to co-sediment with polysomal fractions. Thus, we measured the TP53INP1 protein amount under TNF-α treatment and METTL3 knockdown conditions. TNF-α treatment induced the TP53INP1 protein amount as expected (Figure 5A). Interestingly, METTL3 knockdown led to a more prominent increase in the TP53INP1 amount, parallel to the increase in the rate of apoptosis under this condition (Figure 3B). There appears to be a correlation between the METTL3-mediated modulation of TP53INP1 and the rate of TNF-α-mediated apoptosis. This observation appears to be specific to HeLa cells as we could not confirm it in ME-180 cells (Figure 5B), another human cervix cancer cell line. Parallel to the increase in the TP53INP1 protein amount under TNF-α alone or TNF-α treatment and METTL3 knockdown conditions, we observed an increase in the amount cleaved caspase 3 and 8 (Figure 5C) in HeLa cells, further confirming be the correlation between the METTL3-mediated modulation of TP53INP1 and the rate of TNF-α-mediated apoptosis. Since METTL3 knockdown leads to a further elevation in the amount of TP53INP1 upon TNF-α treatment (Figure 5A), we hypothesized that overexpression of METTL3 should cause a reduction in TP53INP1 amount. As expected, METTL3 overexpression suppressed TP53INP1 amount in TNF-α-treated HeLa cells (Figure 5D). Additionally, METTL3 overexpression improved the survival rate of HeLa cells (Figure 5E).

**Figure 5.**
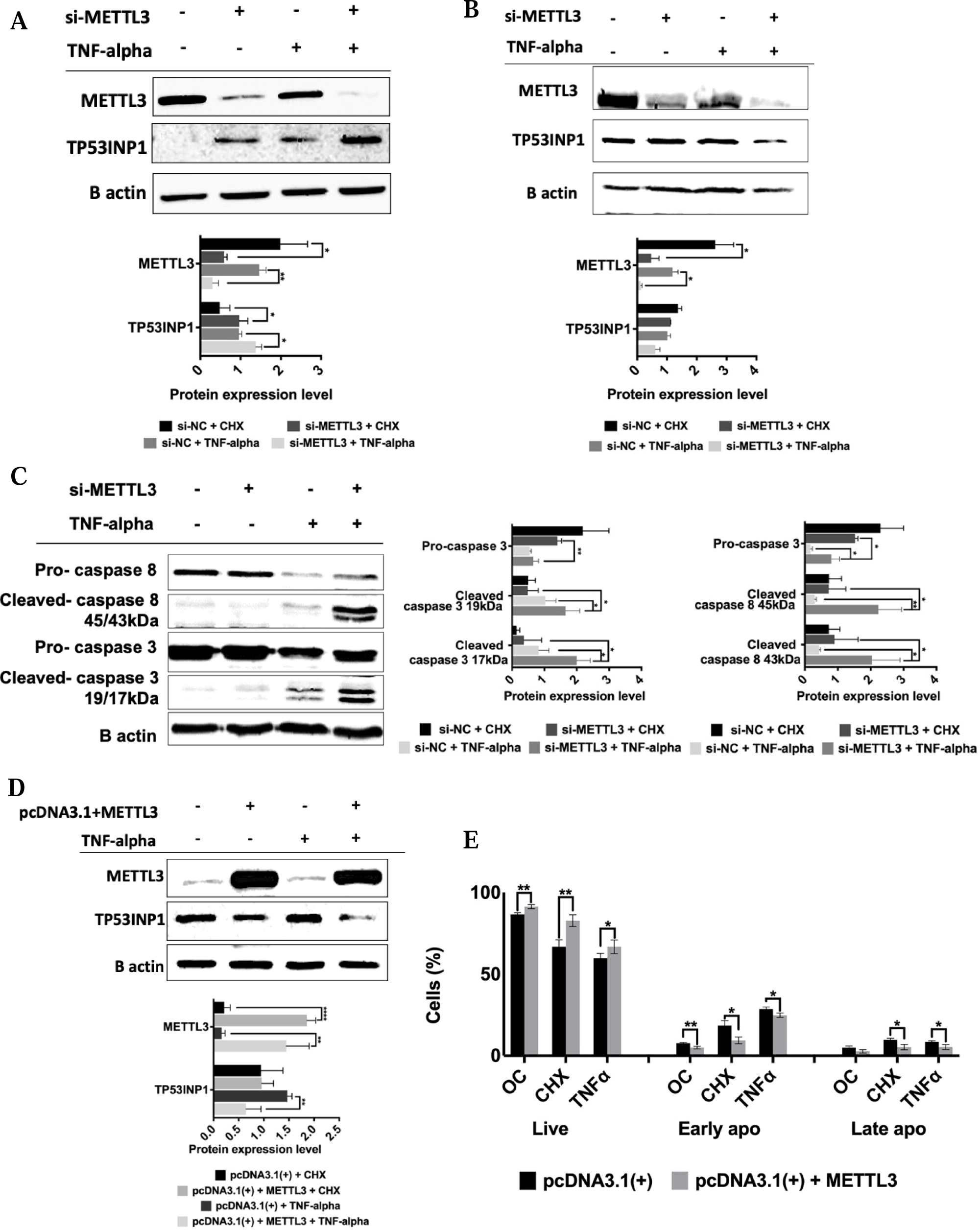
Western blot analyses of METTL3 knockdown cells treated with TNF-α. Cells were transfected with 25 nM control si-NC or si-METTL3 for 72h and incubated with 2.5 ug/mL CHX or 37.5 ng/ml TNF-α for 24h. Western blot assays were performed to examine TP53INP1 expression in METTL3 knockdown and/or TNF-α-treated HeLa (A) and ME180 (B) cells. Total cellular extracts were also assayed for caspase-3 and 8 amounts (C). D. Western blot assays were performed with crude extracts obtained from HeLa cells transfected with the control pcDNA3.1(+) or pcDNA3.1(+) + METTL3 plasmids and treated with 37.5 ng/ml TNF-α. For all western blot experiments, equal amounts of total proteins (25 μg/lane) were fractionated through a 10% SDS-PAGE. Band intensities were normalized against β-actin used as a loading control. n = 3. Two-tailed Student’s t test was performed to determine the statistical significance among groups. Data presented as mean ± SD. *: p≤0.05, **: p≤0.01.

It is highly interesting that *TP53INP1* translation is modulated under the TNF-α treatment condition in a METTL3-dependent manner (Figure 5A). *TP53INP1* is a stress-induced target of p53 (Tomasini et al. 2003), which serves as a proapoptotic tumour suppressor gene. Since TNF-α induces both caspase 8 and 9 cleavage (Figure 1C), we interrogated whether *TP53INP1* translation is also modulated in METTL3 knockdown cells by CP, which induces apoptosis through mitochondria and DNA damage (Florea and Büsselberg 2011). The transcript abundance did not change under CP treatment alone, METTL3 knockdown alone or combined CP treatment and METTL3 knockdown conditions (Figure 6A). We did not detect any change in the polysomal association of TP53INP1 in the control DMSO (Figure 6B) or CP-treated METTL3 knockdown HeLa cells (Figure 6C). Additionally, we did not observe any difference in the protein amount of TP53INP1 in the control DMSO or CP-treated METTL3 knockdown HeLa cells (Figure 6D), clearly suggesting that CP does not modulate *TP53INP1* transcript abundance or polysome association in the control or METTL3 knockdown HeLa cells. These results suggest that METTL3-dependent polysomal association and increase in its protein amount is specific to TNF-α-mediated apoptosis in HeLa cells.

**Figure 6.**
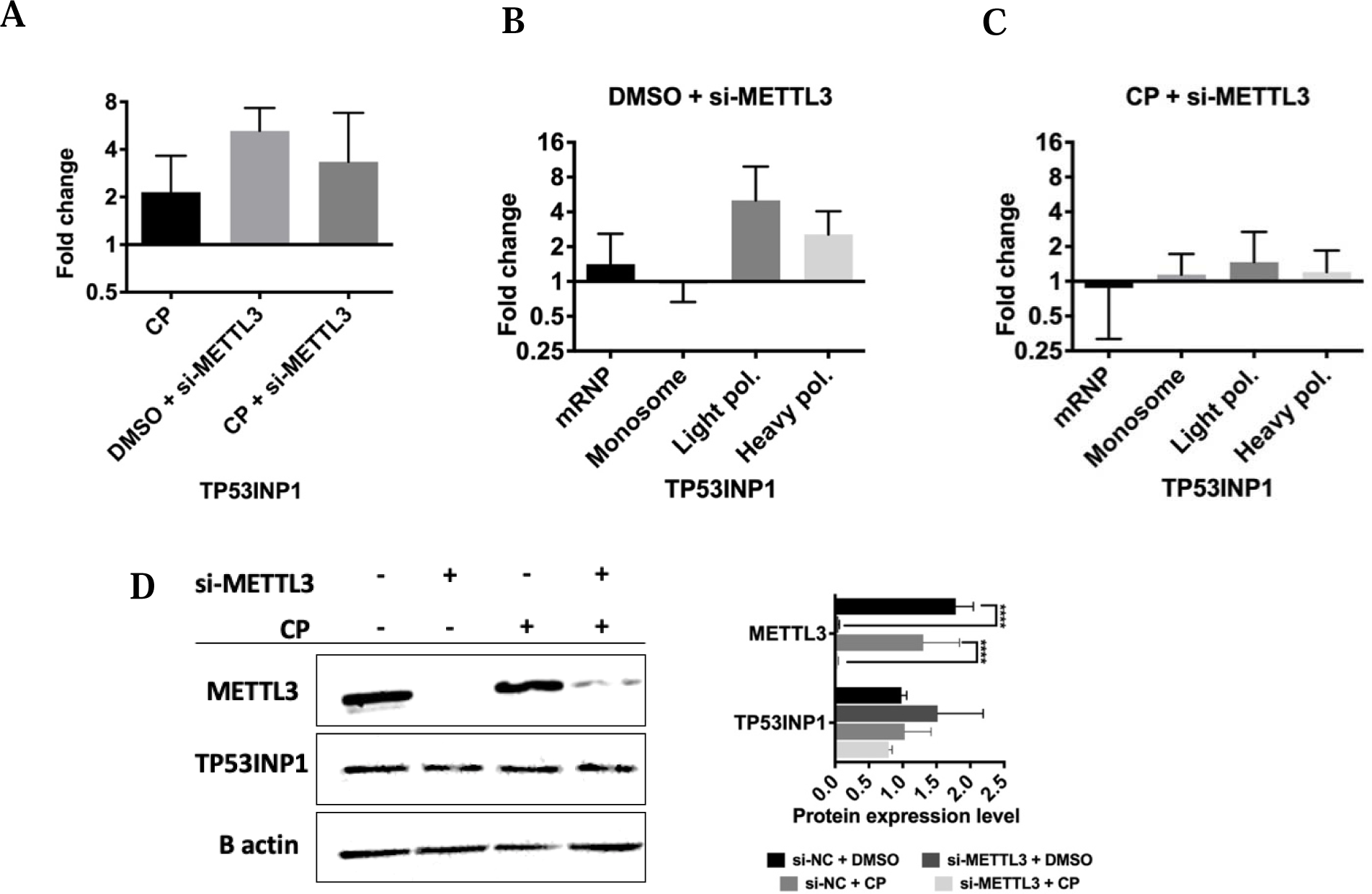
TP53INP1 expression under cisplatin-mediated apoptotic conditions in METTL3 knockdown HeLa cells. A. qRT-PCR analysis of TP53INP1 mRNA abundance in control DMSO (0.1%) or cisplatin-treated (40 µM, 16h) HeLa cells transfected with 25 nM si-METTL3 for 72h. Polysome profile analyses of siMETTL3-transfected HeLa cells treated with control DMSO (0.1%) (B) or cisplatin-treated (40 µM, 16h) (C). Fractionations and qPCR analyses in B-C were performed essentially as described in Figure 5. D. Western blot assay of TP53INP1 protein expression in METTL3 knockdown HeLa cells treated with 40 µM cisplatin (16h). CP, cisplatin. Two-tailed Student’s t test was performed to determine the statistical significance among groups. n =3 biological replicates. Data presented as mean ± SD, *: p≤0.05, **: p≤0.01, ***: p≤0.001****: p≤0.0001.

### METTL3-dependent Modulation of Apoptosis in a Zebrafish Xenograft Model

To assess whether METTL3 inhibition in HeLa cells exerts similar effects *in vivo*, we further evaluated cell death by apoptosis in the larval zebrafish xenografts. Whole-mount immunofluorescence and confocal imaging quantification of DiL+, DAPI+ HeLa cells revealed that neither METTL3 knockdown nor TNF-α treatment alone caused a significant alteration in their apoptosis, indicated by the percentage of cleaved-caspase3 positive cells (Figure 7A-B). In contrast, TNF-α treatment significantly increased apoptotic death of METTL3 knockdown HeLa cells as compared to both TNF-α- or CHX-treated cells treated with control si-NC. These results further support the synergistic action of METTL3 inhibition and TNF-α stimulation on apoptosis induction in HeLa cells.

**Figure 7.**
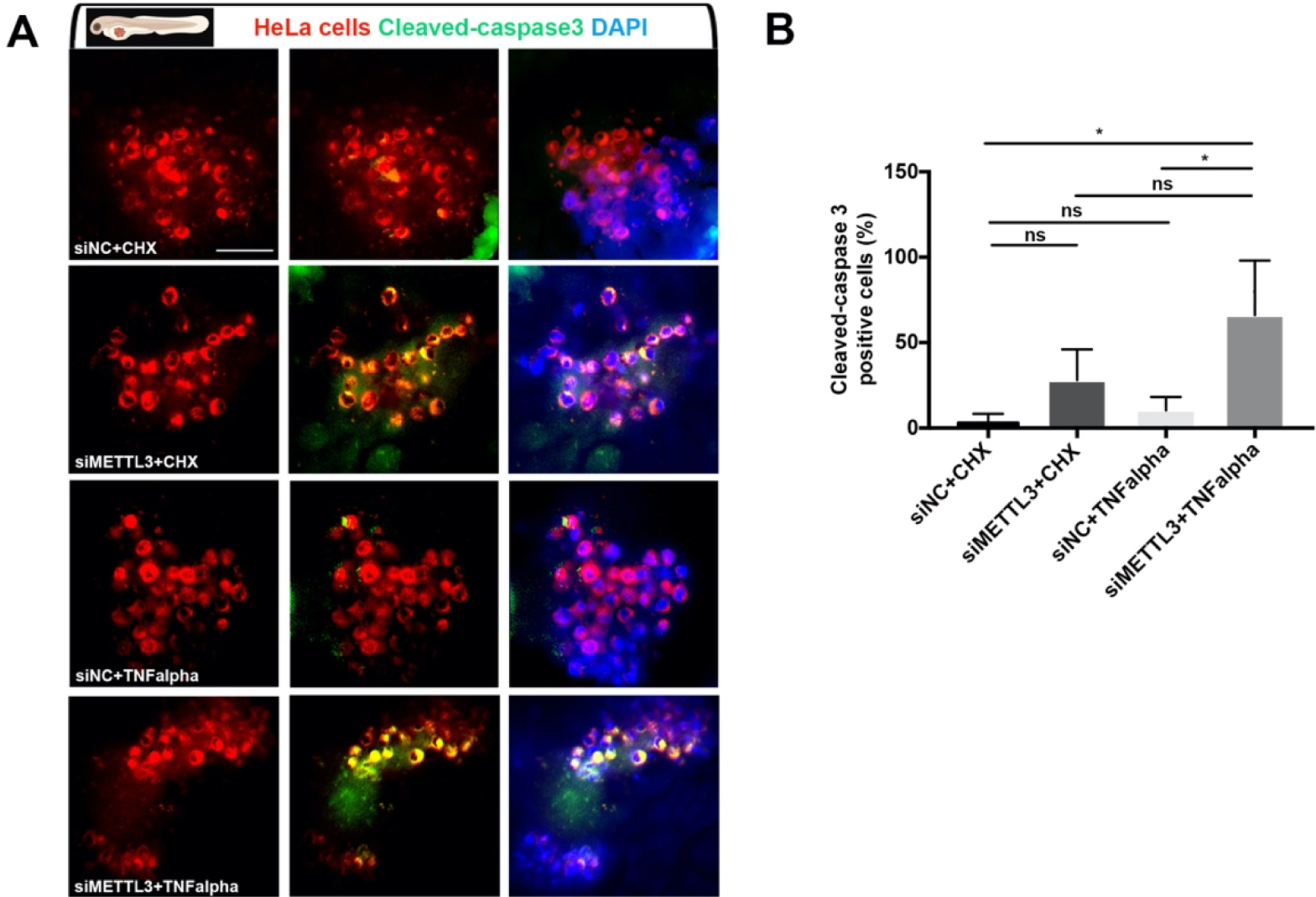
Knockdown of METTL3 synergizes with TNF-α to promote apoptosis of HeLa cells *in vivo*. (A) Representative confocal microscope images of anti-cleaved-caspase-3 (green) staining of 3 dpf (1 dpi) zebrafish larvae xenografted with HeLa cells (red) at 2 dpf, treated with negative control siRNA (si-NC)+CHX, si-METTL3+CHX, si-NC+TNF-α or si-METTL3+TNF-α. (B) Graph showing the percentage of cleaved-caspase3 positive cells in each treatment. Bars represent the average percentage of apoptotic cells counted in each z-stack slice divided by the number of DiL+, DAPI+ nuclei in (A). Larvae were counterstained for DAPI. Scale bars 50μm.

## Discussion

TNF-α is a ligand that modulates numerous cellular phenotypes ranging from cell survival to cell death, especially in the immune system (Webster and Vucic 2020). When coupled with CHX, TNF-α induces apoptosis in HeLa cells by activating the extrinsic apoptotic pathway (L. Wang, Du, and Wang 2008, Figure 1). We provide evidence that the m^6^A RNA methylation machinery is regulated under TNF-α-mediated apoptotic conditions in HeLa cells (Figure 1). Our unbiased approach reveals a plethora of m^6^A RNA marks that might be involved in the TNF-α signalling. Interestingly, *TP53INP1*, a target of p53 known to regulate the intrinsic apoptotic pathway, appears to be modulated translationally by METTL3 either directly or through epitransciptomic modification under TNF-α treatment conditions.

Current studies show that the m^6^A RNA methylation machinery modulates not only cell death and survival but also drug resistance (Dominissini et al. 2012; Zaccara, Ries, and Jaffrey 2019; Xiaowei Zhang et al. 2021). Although METTL3 knockdown leads to cell death in HepG2 cells (Dominissini et al. 2012), we detected a marginal increase in the rate of Annexin-V-positive cells in HeLa cells in the control cells (Figure 3B). However, we observed a marked increase (23.8%) in the rate of apoptosis in the presence of TNF-α (Figure 3B), suggesting sensitization by METTL3 of HeLa cells to TNF-α-mediated apoptosis. qPCR and western blot analyses revealed changes in the expression of writers (Figure 1). Similarly, treatment of HeLa cells with CP downregulates the expression of METTL3 and METTL14 (Alasar et al. 2022), suggesting that downregulation of writer proteins may be a common feature in apoptotic cells independent of the inducers.

We previously reported the complete m^6^A methylome of HeLa cells treated with CP, a universal inducer of the intrinsic apoptotic pathway (Alasar et al. 2022). CP treatment caused differential m^6^A RNA methylation of 132 transcripts associated with apoptosis. On the other hand, in this study, we report that TNF-α treatment leads to perturbations in the m^6^A RNA methylation of 99 apoptotic transcripts. Comparison of the methylation patterns of transcripts under TNF-α and CP treatment conditions revealed that the m^6^A RNA methylome pattern under TNF-α-mediated apoptotic conditions is quite distinct (Figure 2A-C), suggesting the deposition of m^6^A RNA marks in a pathway specific manner. This notion is supported by the observation that the enhanced translation of *TP53INP1* under METTL3 knockdown conditions is attained by TNF-α treatment but not with CP treatment (Figure 5A and 6D). It is currently unknown how the pathway specificity is attained. One can speculate that an apoptotic pathway-specific modulator protein in the writer complex could orchestrate the transcript repertoire of METTL3 by modulating the target specificity. It is also possible that the same m^6^A RNA mark may be recognized differentially by readers in a pathway-specific manner, resulting in a pathway-specific RNA fate.

It remains to be resolved how the apoptotic signalling pathway is connected to the m^6^A RNA methylation machinery. However, once activated, m^6^A marks have a major impact on the fate of mRNAs (Zaccara, Ries, and Jaffrey 2019). The impact of TNF-α treatment on transcript fate was quite different compared to that of METTL3 knockdown. For example, TNF-α treatment perturbed the abundance of candidate transcripts such as *PHLDA1*, *BMF* and *BCL2A1* (Figure 3C). On the other hand, METTL3 knockdown did not cause any change in the abundance of these transcripts, indicating that the abundance of candidate transcripts is probably not modulated by m^6^A RNA methylation under control conditions. Interestingly, under METTL3 knockdown conditions, the abundance of candidate transcripts was highly similar under control and TNF-α treatment conditions. On the contrary, we detected changes in the translational efficiencies of a few of the candidates, *TP53INP1* being the most prominent one (Supplementary Figure 2J, Figure 4J-K, Figure 8). *TP53INP1* is a p53 inducible tumor suppressor gene that activates apoptosis in a caspase-dependent manner (Gironella et al. 2007; Okamura et al. 2001). As part of a positive feedback loop, *TP53INP1* also leads to phosphorylation by HIPK2 and PKCδ of p53 (Tomasini et al. 2003; Yoshida, Liu, and Miki 2006). Since the wild-type p53 is degraded by E6 protein from human papillovirus type 18 in HeLa cells (Scheffner et al. 1990), *TP53INP1* should be activated by p53-independent mechanisms. It is very interesting that *TP53INP1* is activated by TNF-α in HeLa cells (Figure 5).

**Figure 8.**
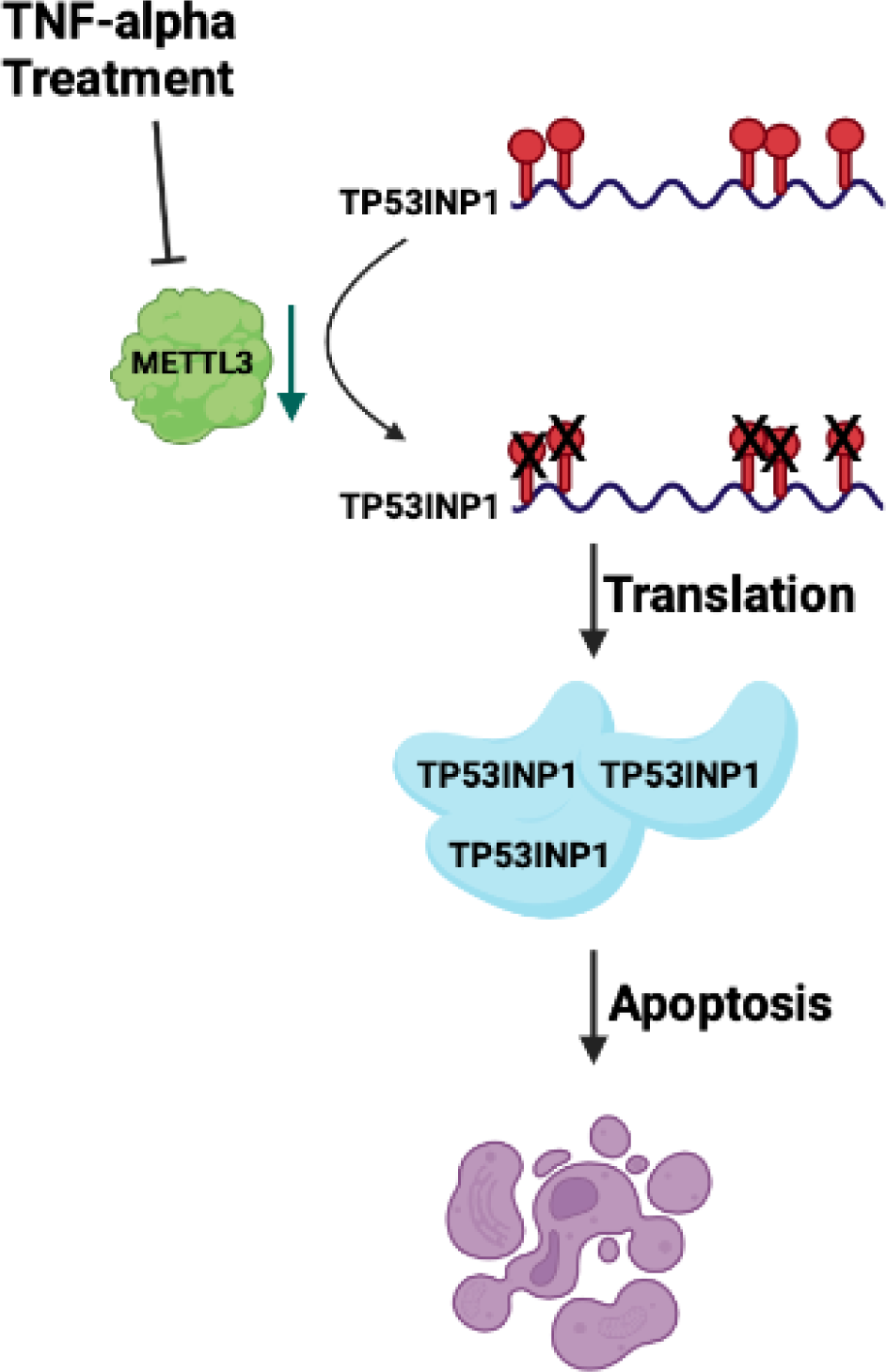
Schematic representative of model.

We report the TNF-α-METTL3-TP53INP1 axis as a new mechanism that appears to modulate the extrinsic pathway of apoptosis in HeLa cells. We detected five adenosine residues which are methylated under control conditions (Figure 8). Although there appears to be a nice correlation between the enhanced translation of *TP53INP1* transcript and its reduced m^6^A RNA methylation under TNF-α treatment or METTL3 knockdown conditions (Figure 8), we cannot ascertain which particular methylation mark is responsible for this translational enhancement. Further studies are required to delineate the contribution of each m^6^A mark on *TP53INP1* translational efficiency. For example, RNA-guided RNA interfering activity of chimeric Cas13 proteins (dCas13-METTL3 or dCas13-FTO) could be highly useful for this purpose.

## Acknowledgements

The authors would like to thank İpek Erdoğan Vatansever, Özgür Okur and Murat Delman for flow cytometry analyses, and Biotechnology and Bioengineering Application and Research Centre (IZTECH, Turkey) for the instrumental help. This study is funded by the Scientific and Technological Research Council of Turkey (TUBITAK Project No: 217Z234 to BA). GO Lab is funded by EMBO Installation Grant (IG 3024).

## Author Contributions

BA contemplated the project. ÖT performed polysome experiments. UC, EI and GO performed zebrafish xenograft experiments. MA constructed the heat maps. AAA, BS and YG performed all other experiments. AAA, GO and BA wrote the manuscript, all authors read and approved the manuscript.

## Conflict of Interest

The authors declare that they have no conflict of interest.

## Ethics Statement

The animal study was reviewed and approved by the Animal Experiments Local Ethics Committee of Izmir Biomedicine and Genome Center (IBG-AELEC).

## Notes

### Competing Interest Statement

The authors have declared no competing interest.

### Summary of Updates

Typing errors have been fixed without any changes in the data or text.

