## Supplemental Figures 1-3 for "Epitranscriptomics m^6^A analyses reveal distinct m^6^A marks under tumor necrosis factor α (TNF-α)-induced apoptotic conditions in HeLa cells" (from another version of this preprint: version 1, 2023-05-20)

**Alasar et al. Supplementary Figure 1**

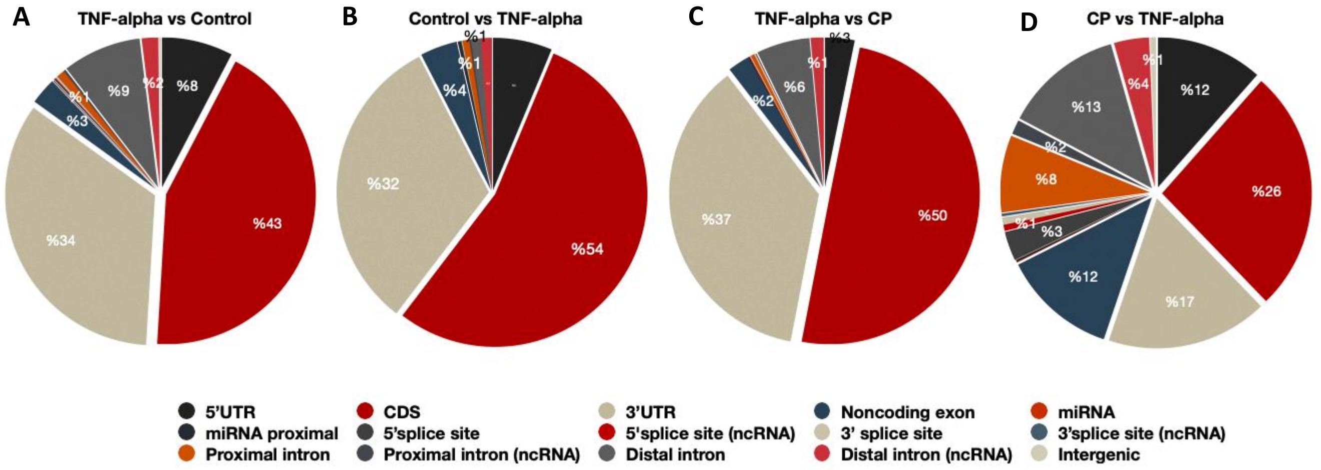


**Supplementary Figure 1. m$^6$A analysis by miCLIP in TNF-α /CP treated HeLa cells.** Pie chart depicting transcripts upregulated (A) or downregulated (B) in m$^6$A methylation in TNF-α-treated HeLa cells, upregulated (C) and downregulated (D) m$^6$A modification in TNF-α induced apoptosis compared to CP treatment. CP, cisplatin.

**Alasar et al. Supplementary Figure 2**

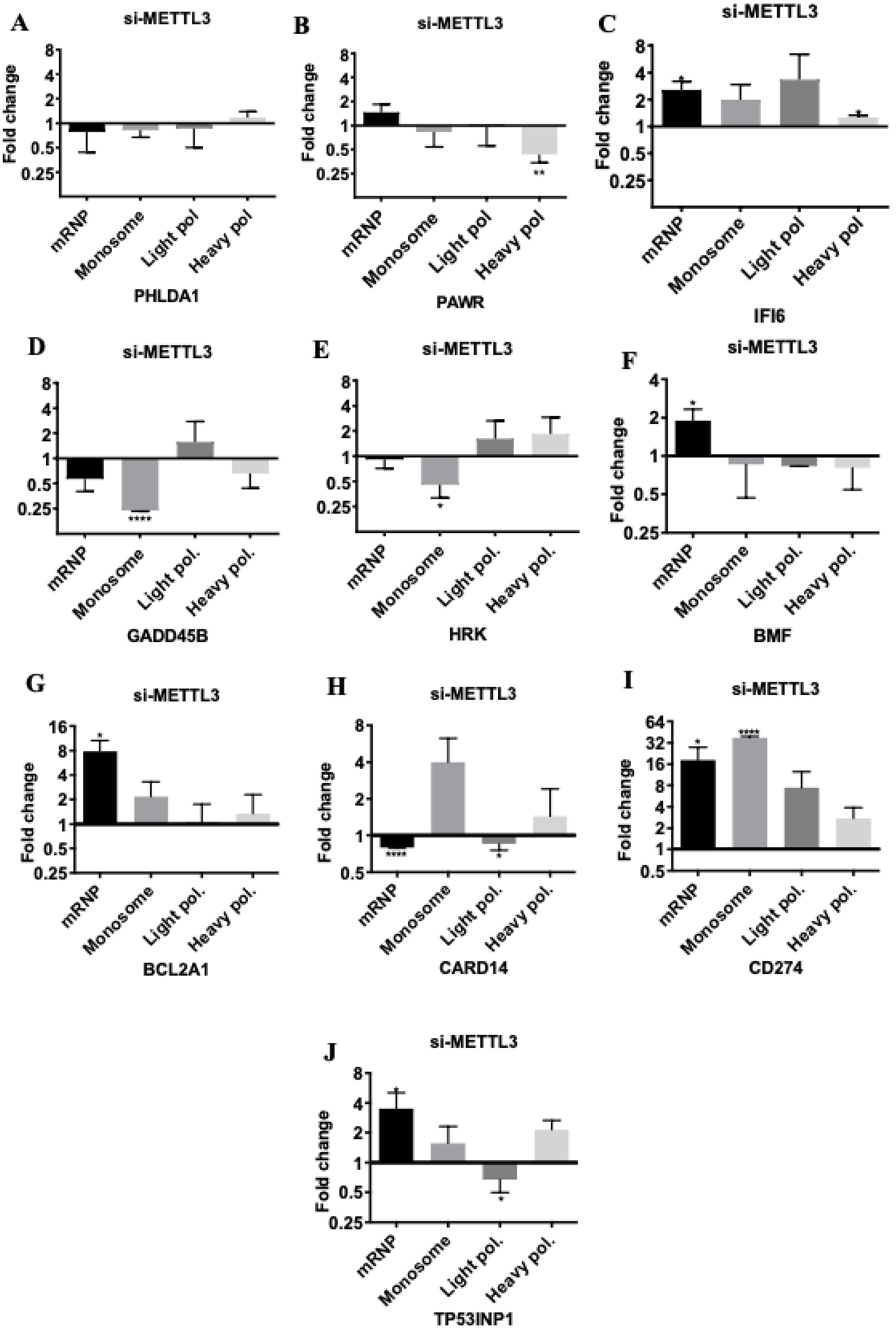

**Supplementary Figure 2. Effects of METTL3 knockdown on translational efficiencies of candidate mRNAs.** qPCR analyses of *PHLDA1* (A), *PAWR* (B), *IFI6* (C), *GADD45B* (D), *HRK* (E), *BMF* (F), *BCL2A1* (G), *CARD14* (H), *CD274* (I) and *TP53INP1* (J) in HeLa cells upon METTL3 knockdown. HeLa cells transfected with 25 nM si-NC (control) or si-METTL3 for 72h were subjected to polysome profiling as explained in Materials and Methods. Four fractions, namely mRNP, monosome, light and heavy polysomes, were collected based on $A_{254}$ reading. GAPDH was used for normalization. All experiments were conducted in triplicates. Data are presented as mean values ± SD. Two-tailed Student's t test was performed to determine the statistical significance among groups. **: p≤0.05, **: p≤0.01, ****: p≤0.0001.

**Alasar et al. Supplementary Figure 3**

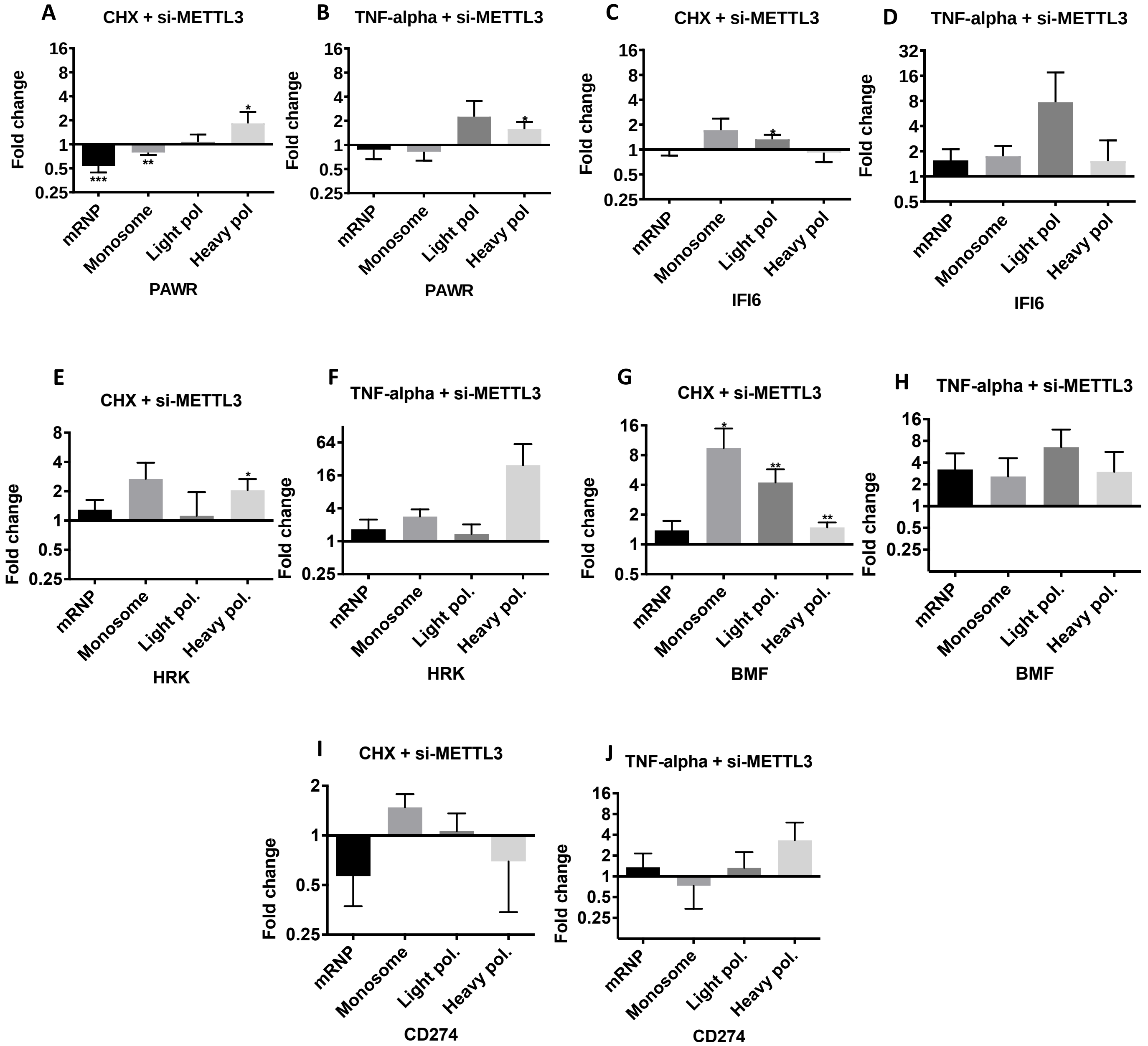


**Supplementary Figure 3. Effects of METTL3 knockdown and TNF-α treatment on translational efficiencies of candidate mRNAs.** mRNA abundance of *PAWR* (A, B), *IFI6* (C, D), *HRK* (E, F), *BMF* (G, H) and *CD274* (I, J) mRNAs in mRNP, monosome, light polysome and heavy polysome fractions was quantified by qPCR in TNF- α treated METTL3 KD HeLa cells. n = 3 biological replicates. Two-tailed Student's t test was performed to determine the statistical significance among groups. Data presented as mean ± SD, *: p≤0.05, **: p≤0.01, ***: p≤0.001.
